# Intraflagellar transport trains can turn around without the ciliary tip complex

**DOI:** 10.1101/2021.03.19.436138

**Authors:** Adrian Pascal Nievergelt, Ilia Zykov, Dennis Diener, Tim-Oliver Buchholz, Markus Delling, Stefan Diez, Florian Jug, Ludĕk S̆tĕpánek, Gaia Pigino

## Abstract

Cilia and flagella are microtubule doublet based organelles found across the eukaryotic tree of life. Their very high aspect ratio and crowded interior are unfavourable to diffusive transport for their assembly and maintenance. Instead, a highly dynamic system of intraflagellar transport (IFT) trains moves rapidly up and down the cilium. However, the mechanism of how these trains turn around upon reaching the ciliary tip has remained elusive. It has been hypothesized that there exists a dedicated calcium-dependent protein-based machinery at the ciliary tip to mediate this conversion. In this work, we use physical and chemical methods to manipulate IFT in the cilia of the unicellular green alga *Chlamydomonas reinhardtii* to show that no such stationary tip-machinery is required for IFT turnaround. Instead, we demonstrate that the conversion from anterograde to retrograde IFT trains is a calcium independent intrinsic ability of the IFT system.

## Introduction

Motile cilia and eukaryotic flagella are highly ordered organelles that serve a variety of vital functions such as single-cell locomotion, left-right symmetry breaking in mammalian development, or chemomechanical sensing (***Shah et al., 2009***). The cytoskeleton of a cilium is the axoneme, a nine-fold symmetric structure consisting of nine microtubule doublets, which are highly decorated with a plethora of motor and regulatory complexes, arranged around two central microtubule singlets known as the central pair (***Nicastro et al., 2006***). The axonemal structure is highly conserved across the eukaryotic phylogeny. With an aspect ratio of typically 30–60, the assembly and maintenance of motile cilia is physically infeasible by diffusive transport alone. Instead, the material involved in assembly, disassembly and signaling is transported along the cilium by an active multi-protein-complex transport machinery referred to as intraflagellar transport (IFT).

IFT was first discovered in the cilia/flagella of the unicellular green alga *Chlamydomonas reinhardtii*. Using their two cilia, Chlamydomonas can adhere and slowly glide over glass surfaces. During gliding one can observe densities within the cilium that move from the cell body towards the tip (anterograde direction) and back again (retrograde direction) (***Kozminski et al., 1993***). The observed densities have been shown to consist of long chains of IFT particles, which themselves consist of two protein complexes (IFT-A and IFT-B) as well as motors and cargo (***Cole et al., 1998***; ***Pigino et al., 2009***; ***Taschner and Lorentzen, 2016***; ***Jordan et al., 2018***). Since the initial discovery of IFT, fluorescent labeling has allowed a much more detailed observation of IFT particles *in vivo* (***Engel et al., 2009***). Using correlated fluorescence microscopy and electron tomography, we have shown that IFT inherently avoid collisions: anterograde trains, driven by the heterotrimeric motor kinesin-II walk along the B-tubule while retrograde trains, driven by dynein–1b motors, walk along the A-tubule (***Stepanek and Pigino, 2016***). Additionally, we have shown that anterograde and retrograde trains are structurally different (***Stepanek and Pigino, 2016***). Upon reaching the tip of the cilium, the kinesin-II motors dissociate from the IFT trains (***Chien et al., 2017***) and convert to retrograde trains by an unknown mechanism (see fig. 1A).

**Figure 1.**
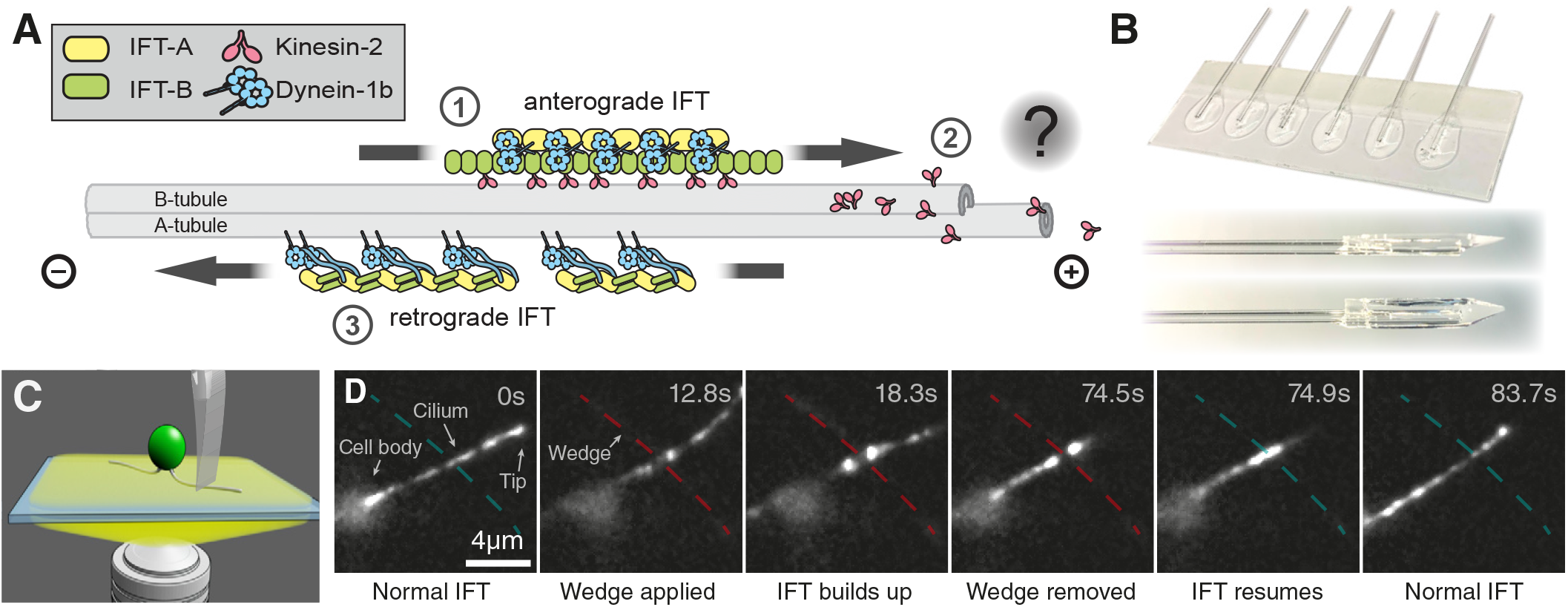
The intraflagellar transport (IFT) machinery of gliding Chlamydomonas can be observed and manipulated in total-internal-reflection microscopy (TIRF). **A)** ➀ Anterograde trains, driven by the heterotrimeric motor kinesin-II move towards the ciliary tip. ➁ At the ciliary tip, kinesin-II dissociates from the train and the trains convert into retrograde form by an unknown mechanism. ➂Reassembled retrograde trains move away from the tip towards the cell body by the dynein-1b motor specific for IFT. **B)** A sharp, but soft wedge is made by drop-casting silicone onto a supporting capillary, followed by trimming off excess after curing. **C)** Illustration of mechanical blockage by lowering a silicone wedge onto the cilium of a *C. reinhardtii* cell by micromanipulator on a TIRF microscope. **D)** Individual frames from a TIRF movie show how IFT is locally and reversibly blocked inside a cilium by the force applied using a silicone wedge (dashed line, non-blocking in blue, blocking in red).

A prominent hypothesis is that there is a dedicated machinery at the ciliary tip which is responsible for the conversion of trains from their anterograde to their retrograde structure. This hypothesis is supported by the observation of large protein complexes that cap the microtubule plus-ends and form the so called ciliary tip complex (***Fisch and Dupuis-Williams, 2011***). Here, we test this hypothesis by directly interfering with IFT by either creating a physical barrier along the cilium, using a newly developed soft silicone wedge technique, or by chemically inhibiting the motion of trains using lithium chloride.

## Results

We first set out to physically block and observe IFT in Chlamydomonas cells. In order to apply a gentle but constant localized force onto a specific spot on the cilia of gliding cells, we fabricated a sharp silicone wedge by drop casting silicone resin onto the end of a glass capillary lying on a glass slide. We then cured the drop on a hot-plate and trimmed away the excess material with a scalpel before lifting the wedge from the glass slide (fig. 1B). Finally, this wedge was mounted on a hydraulic micro-manipulator which was installed on a total internal reflection fluorescence (TIRF) microscope. During TIRF imaging, the wedge was carefully lowered onto the cilium of a gliding cell (see fig. 1C). Insufficient pressure would result in IFT trains passing the partial barrier created by the wedge. Excessive pressure, instead, would damage or even sever the cilium, resulting in an immediate stand-still of all visible IFT trains, presumably due to the leakage of ATP from the cut site. The soft wedge, when carefully controlled by the micro-manipulator, allows fine control over the degree at which IFT trains are blocked: light pressure will sporadically allow trains to squeeze underneath the created barrier, becoming very visible in TIRF as they move closer into its evanescent field. When using just the right amount of pressure, IFT trains can no longer pass through the blockage we created. As a consequence, we observe no fluorescence underneath the wedge and a striking accumulation of trains on both sides of it (see fig. 1D). Once all trains caught between the wedge and the ciliary tip have accumulated at the blockage site, the brightness of the more distal accumulation spot remains constant and the entire distal part of the cilium is depleted of fluorescence. This observation empathizes (*i*) the completeness of the blockage, and (*ii*) that retrograde trains do not spontaneously convert into anterograde trains. Upon removing the wedge, IFT resumed through the previously blocked region without any noticeable residual effect. Even when we performed the experiment multiple times on the same spot, IFT blockage and recovery took place just as described above (see video 1).

**Video 1**. Intraflagellar transport in the cilium of a Chlamydomonas cell is repeatedly blocked and subsequently released by application of a silicone wedge.

To facilitate the analysis of the motion of IFT trains in TIRF time-series, we produced kymograms which display the positions of the trains as a function of time (fig. 2). In kymograms, trains moving at a constant speed appear as straight lines, with anterograde trains (diagonal green traces) easily distinguishable from retrograde trains (diagonal magenta traces) or stalling trains (uncolored horizontal lines). Additionally, the lateral gliding motion of the cell is directly visible from the position of the ciliary tips. As can be seen in fig. 2A, shortly after lowering the wedge onto the cilium, trains start accumulating on both sides of the wedge. The tip side of the cilium (upper part of the kymogram) gets rapidly depleted of trains and retrograde trains accumulate distally with respect to the wedge. Anterograde trains, in contrast, are still injected from the cell body into the cilium and travel up to the barrier where they accumulate. Notably, on the cell body side of the barrier (proximal side), about 2.5 seconds post-blocking, retrograde IFT trains can be observed as they depart from just below the barrier towards the cell (see fig. 2B). Soon after the onset of this phenomenon, the frequency of departing retrograde trains returns to normal. When the wedge is retracted, the motion of the artificially arrested trains on both sides of the wedge resumed to regular antero-grade and retrograde motion. These data clearly show that the conversion from anterograde to retrograde IFT can and does take place in the absence of the ciliary tip.

**Figure 2.**
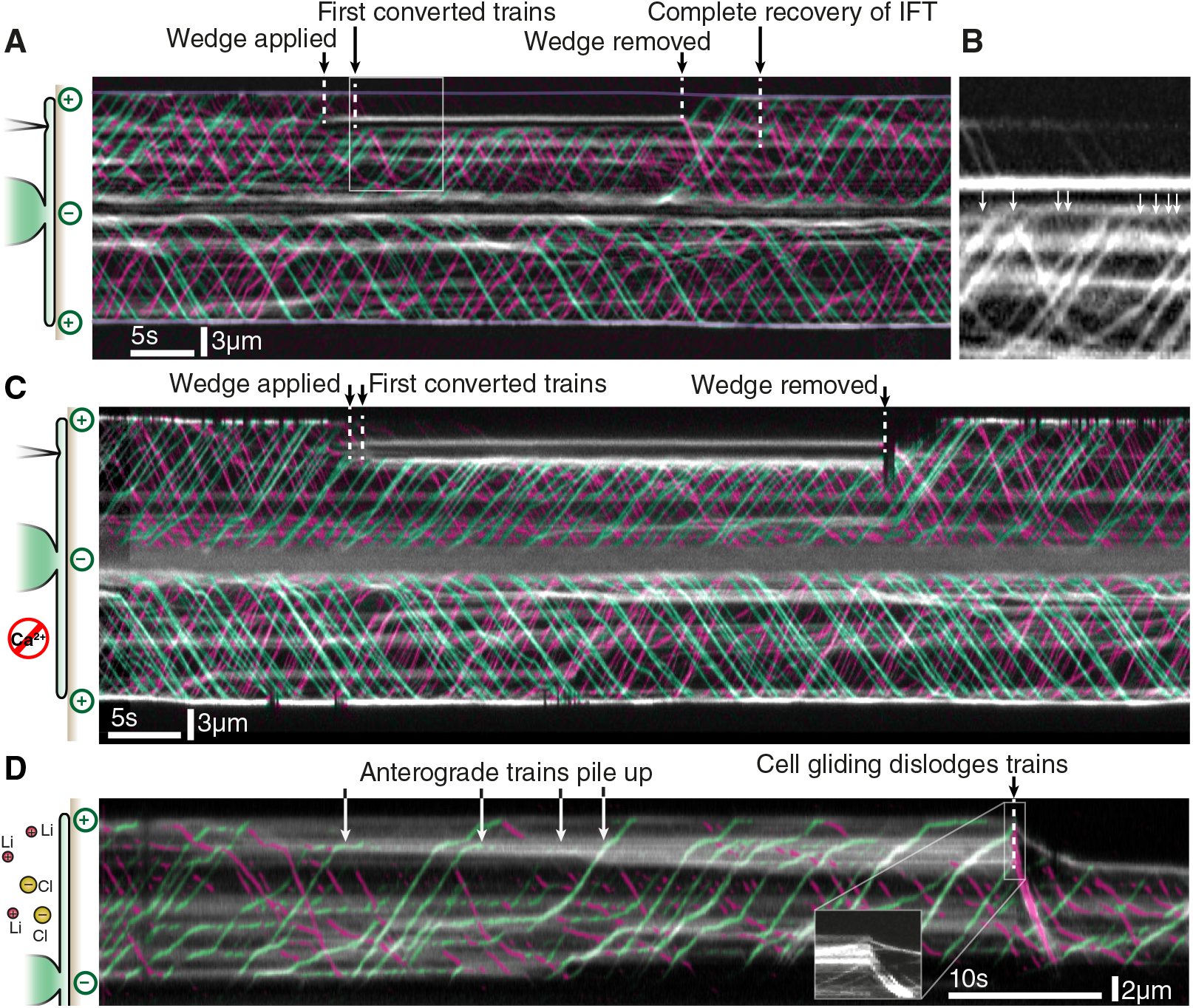
Kymograms visualizing the time-dependent response of the IFT machinery to external manipulation. **A)** Compressing the cilium with a soft wedge creates a small band where IFT can not pass (horizontal white lines in the kymogram, created by the accumulation of stopped fluorescent trains). After the blockage is applied, trains build up and shortly after start moving retrograde. The approximate location of the ciliary tips is outlined for clarity in purple. **B)** Detail view of the region between the cell body and the block in **A)**. About 2.5s after IFT is locally blocked, retrograde trains can be seen moving towards the cell body at normal speed (white arrows). **C)** Kymogram of wedge-blocking experiment in severely calcium-depleted cilia. Transport and conversion show no discernable difference to **A)** where calcium is available. **D)** In lithium-treated cells, the motors are chemically inhibited and pile up at variable distances from the ciliary tip. Gliding of the cell dislodges the stuck trains which immediately move in retrograde direction at high speed (≈ 10 µm s^-1^) (insert). Anterograde trains are colored in green, while retrograde trains are colored in magenta for clarity.

It has been proposed that calcium-dependent kinases could be involved in the regulation of anterograde IFT motor kinesin-II (***Liang et al., 2014***). We reasoned that the application of the wedge could induce a localized tension on the ciliary membrane and potentially open calcium channels locally. If that were the case, the resulting calcium influx from the buffer could activate the calcium-dependent kinases and locally induce the anterograde-to-retrograde conversion. Therefore, to test whether free calcium is a co-factor in the anterograde–to–retrograde conversion at the barrier, we performed the wedge-blocking experiment in calcium-free tris-acetate-phosphate (TAP) medium with ethylene glycol-bis(/3-aminoethyl ether)-N,N,N’,N’-tetraacetic acid (EGTA) added as an additional strong calcium chelator. This medium is further referred to as TAP-Ca. To characterize the effect of this environment, we observed Chlamydomonas cells with a fluorescent free-calcium-reporter (GCaMP6) expressed as a fusion protein to the dynein regulatory complex protein 4 (DRC4). In normal TAP medium, these cells exhibit periodic flashes of free calcium in their cilia, which are also observed when the cilia are only lightly touched by the wedge (see video 2).

**Video 2**. In presence of calcium, the cilia of *Chlamydomonas reinhardtii* cells with a fluorescent free calcium reporter (GCaMP6) exhibit flashes indicating free calcium.

Instead, we observed that in TAP-Ca, cells exhibit a total lack of spontaneous or induced flashes (see video 3), indicating that cilia in such an environment are almost completely depleted of free calcium.

**Video 3**. In completely calcium free medium, there is a complete absence of flashes in the cilia of *Chlamydomonas reinhardtii* cells, indicating a total lack of free calcium in the axoplasm.

When repeating the wedge-blocking experiment in TAP-Ca medium, anterograde IFT trains still convert to retrograde trains and no noticeable differences to the reported observations in regular TAP medium can be observed (see 2C). Therefore, we conclude that calcium is not required for anterograde-to-retrograde conversion.

To complement the results from mechanically blocking trains, we treated cells with 13 mM of LiCl, which inhibits IFT motors (***Wilson and Lefebvre, 2004***), thereby causing frequent pauses in the motion of trains (see 2D). This inhibition of motility results in the accumulation of anterograde IFT trains along the cilium, which are stuck and unable to dislodge on their own. However, we observed that the onset of the gliding motion of a cell is often sufficient to restore the motility of stuck trains. Interestingly, as can be seen in fig. 2D, a stuck trains always reactivated in retrograde direction, even if it was an anterograde train when stalling. Hence, treatment of Chlamydomonas with LiCl also shows anterograde-to-retrograde conversion, providing additional arguments for this process to not require the ciliary tip.

After these experiments, we wondered if anterograde-to-retrograde conversion dynamics differ between their natural appearance at the ciliary tip and the wedge-induced occurrence in our experiments. To this end, we quantified the frequency of retrograde trains forming at *(i)* the ciliary tip after first depleting the tip of all IFT material by wedge-blocking and then releasing anterograde trains by removing the wedge, and at *(ii)* the proximal side of the wedge directly after onset of wedge-blocking. We have manually annotated retrograde trains in fifteen blocking experiments on five different cilia and report our findings in fig. 3.

**Figure 3.**
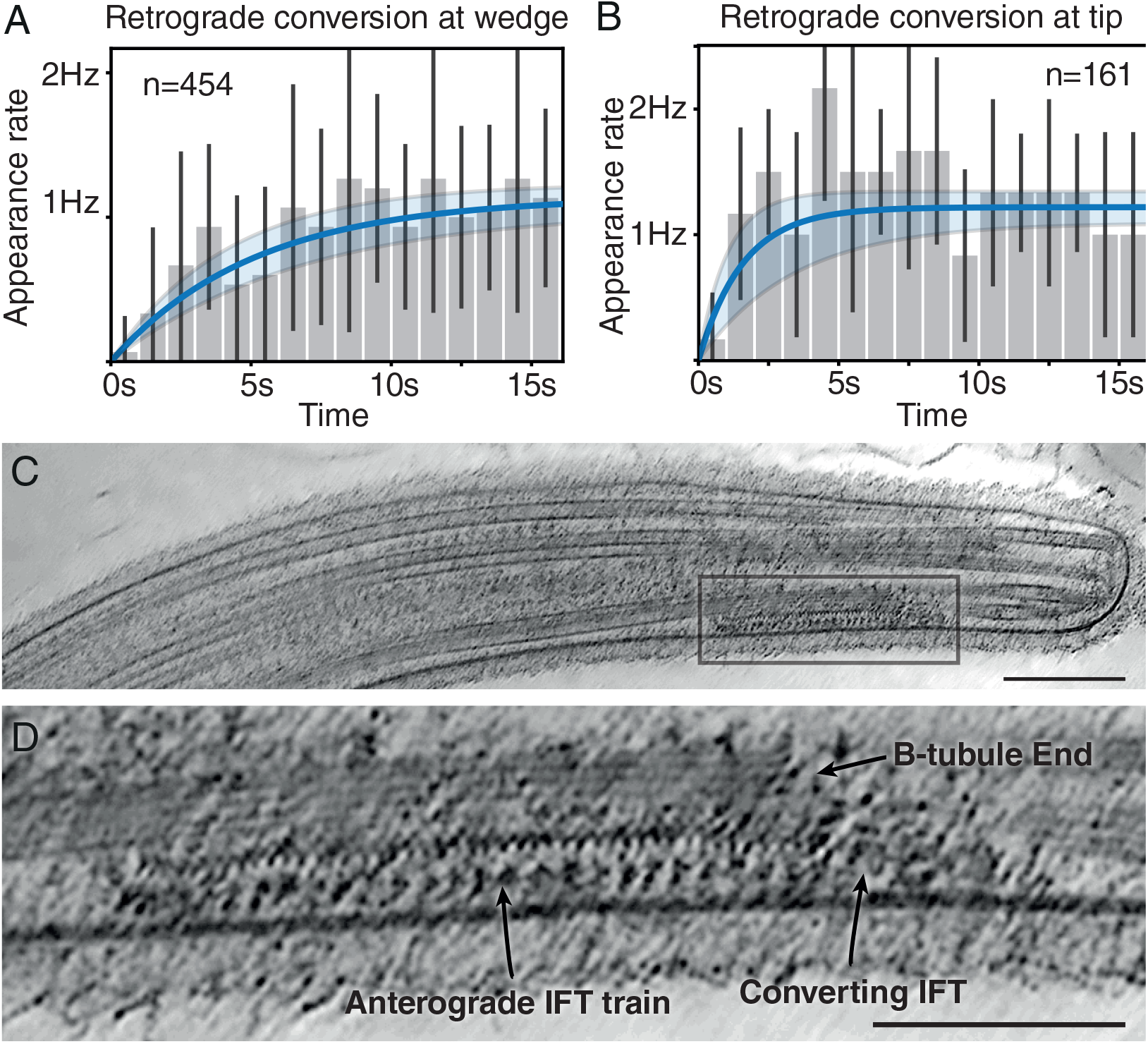
Kinetics of anterograde to retrograde transport conversion of intraflagellar transport trains. **A)** Retrograde train appearance frequency at physical blockage. Transport recovers to normal steady-state level with a time constant of (5.1 ± 1.2) s (first-order response fit in blue, uncertainty bounds in light gray). **B)** Retrograde train appearance frequency at the ciliary tip after releasing the physical blockage. Steady-state retrograde transport recovers with a time constant of (1.60 ± 0.84) s (fit same as A). **C)** Tomographic slice through ciliary tip showing an anterograde train in the process of disassembly (scale bar 100nm). Highlighted rectangular region is magnified in **D)**, showing loss of structural cohesion of the train localized at the end of the B-tubule (scale bar 100nm).

Figure 3A shows the rate at which retrograde trains depart from the wedge area after blocking transport. While very rapidly converting trains are occasionally observed, the conversion of the majority of the trains requires several seconds. Assuming the conversion frequency can be modeled as a first-order system (fitted blue line with light gray uncertainty bounds), we find that IFT returns to its nominal conversion frequency with a time constant of (5.1 ± 1.2) s after blocking. In order to compare this to natural conversion at the tip, we have repeated the same annotation and analysis for the departure times of retrograde trains at the IFT-depleted tip. Since transport is in each experiment blocked at a different distance from the tip, we measured departure times relative to the first arriving anterograde train. 3B shows the analysis of six such experiments on three different cilia. In contrast to conversion at the wedge, IFT returns to the steady-state about three times faster, with a time constant of (1.60 ± 0.84) s. These values are in-line with previous reports in the literature (***Chien et al., 2017***). Hence, while anterograde-to-retrograde conversion can happen in the absence of the ciliary tip, our findings also show that the rate of conversion is higher at the tip than it is when induced by our wedge-experiments.

In order to find trains that are in the process of anterograde-to-retrograde conversion, we performed cryo-electron tomography on intact ciliary tips (see figure 3C). In fig. 3D we show a cry-oCARE (***Buchholz et al., 2019b***) denoised tomographic slice through a train located in close proximity to the distal end of the axoneme. While we can currently not point at underlying molecular mechanisms, our data offers interesting structural observations into this conversion process. Most importantly, such trains show the expected repeating densities of anterograde trains (***Jordan et al., 2018***) as long as they are engaged with the B-tubule of the microtubule doublet they are walking on (***Stepanek and Pigino, 2016***). This structure is rapidly lost after trains disengage from the B-tubule, potentially pointing at a simple explanation why conversion rates are faster at the ciliary tip, where microtubules end and trains can easily disengage by simply walking off their rails.

## Discussion

In this work we demonstrate that IFT trains can reverse their directions in the absence of the ciliary tip. When mechanically blocked with an easily made device, anterograde trains spontaneously convert into retrograde trains. We observe a short delay before reorganized trains start returning to the cell body at a rate which is comparable to the unperturbed case at the ciliary tip. Additionally we showed that the exact same behaviour can be observed in cilia that are calcium-depleted, indicating that this conversion process is not calcium-dependent. Finally, by observing chemically inhibited trains, we offer another way by which anterograde-to-retrograde IFT train conversion can be induced at locations far away from the ciliary tip. We assume that the chemical inhibition of motors causes a different form of mechanical blockage – one that does not require an external force to be applied.

One could argue that anterograde trains which accumulate at the blockage might carry cargo that, upon cessation of motion, assemble a tip–complex-like machinery which then mediates the train conversion. However, when the blockage is released, we never observed additional trains to be stopped and train conversions instantly seizes.

Our data indicate that the conversion from anterograde to retrograde transport is an intrinsic property of IFT trains with no dedicated tip machinery required to enable this process. Even though the ciliary tip is not necessary for conversion, this process is roughly three times faster at the tip than it is at a wedge-blockage site. Based on this kinetic difference and our cryo-tomography data, we propose a reorganization-after-derailment model, where structural reorganization of anterograde trains require the disengagement from the B-tubule of the microtubule doublet they are walking along. While anterograde trains exhibit a well known and highly regular conformation when travelling along the B-tubule (***Jordan et al., 2018***), upon loss of contact with the microtubule, either by moving past the microtubule end or by piling up and disengaging at a blockage site, these trains lose their conformation and undergo noticeable structural changes before forming functional retrograde trains.

Our work is an important step towards understanding the intricate molecular mechanisms that underlie the intraflagellar transport – a system relevant for all cilia-mediated functions.

## Materials and Methods

### Chlamydomonas cell culture

IFT54-mNeon-IFT139-mCherry cells were a kind gift from Karl Lechtreck, University of Georgia. The cells were inoculated from TAP–agar plates into TAP medium and grown in aeration bottles connected to clean air under circadian conditions to early-to mid-log phase. Cells for wedge-experiments were harvested at 1000×g for 10 min and subsequently resuspended in TAP medium supplemented with 1 mM EGTA (ethylene glycol-bis(*β*-aminoethyl ether)-N,N,N’,N’-tetraacetic acid, E3889, Sigma Aldrich, USA). Cells for LiCl experiments were likewise washed in just TAP buffer.

### Transgenic strain creation

#### IFT46-mNeonGreen

The EcoRI-EcoRV YFP containing fragment in pE345 (IFT46-YFP under control of PSAD promoter and IFT46 5’ UTR and unknown 3’UTR; contains IFT46 genomic DNA; PAR resistance) was replaced by an EcoRI-EcoRV fragment encoding the mNeonGreen ORF. The fragment was codon optimized for expression in Chlamydomonas. Codon optimization was done with GenSmartTM Codon optimization tool (GenScript) and synthesized by Integrated DNA Technology. The resulting plasmid was linearized with PsiI.

CC-4375 *ift46-1::NIT1* mt+ cells were obtained from the Chlamydomonas Resource Center and grown in TAP to a dense colony. Cells were havested at 800RCF and subsequently had their cell walls removed with autolysin. 400uL of concentrated cells were then mixed with 600 ng of linearized plasmid and vortexed with 0.5mm Zirconia/Glass beads (11079105z, BioSpecProducts, Bartlesville, OK, United States) for 30s before plating on TAP agar plates with 10 µg mL^-1^ Paromomycin for positive selection. Colonies were subsequently analyzed by western blotting for IFT46.

### PF2-GcAMP6

The PF2-GFP plasmid obtained from Mary Porter as a gracious gift. The chlamydomonas PF2 (DRC4) was amplified by PCR using the PF2-GFP plasmid as a template and cloned into pBR25 using XhoI and BamHI sites. GCaMP6 was amplified using PCR and cloned in-frame at the unique BamHI at the c-terminal of PF2.

The paralyzed flagella strain *pf2* (cc1025, Chlamydomonas Resource Center), was cotransformed with 1 µg of PF2::pGCaM6 and 1 µg pSI103, encoding an aphVIII paramomycin resistance cassette (***Sizova et al., 2001***), using the glass bead method (***Kindle, 1990***), but without the addition of polyethylene glycol. Following transformation the cells were plated on TAP agar plates containing 10 µg/ml paramomycin and resulting colonies were screened for motility after transfer to TAP medium in microtiter wells. Flagella of swimming cells were screened by western blotting with anti GFP for the PF2::pGCaM6.

### Wedge preparation

Glass capillaries (BF100–50–7.5, Sutter instruments, USA) were placed on microscope slides with double-sided adhesive tape and a drop of polydimethylsiloxane (PDMS, Sylgard 184, Dow Corning, USA), premixed at 10:1 monomer-crosslinker ratio, applied to the end of the capillary. The slice was subsequently places on a hot-plate at 120 °C to 140 °C to cure the PDMS for several minutes. Finally, the sides of the PDMS drop were trimmed off with a scalpel and the device carefully delaminated from the microscope slide right before use.

### TAP-Ca medium preparation

Calcium-free TAP-Ca medium was prepared by dissolving 2.42 g of Tris-Base (93362, Merck KGaA, Darmstadt, Germany) to 850 mL deionized water. 25mL of TAP-Ca salts (15 g L^-1^ NH_4_Cl, 4 g L^-1^ MgSO_4_, 2 g L^-1^ KCl) and 1mL of Phosphate solution (288 g L^-1^ K_2_HPO_4_, 2 g L^-1^ KH_2_PO_4_) and 1 mL of Hutner’s trace elements (Chlamydomonas resource center) were added sequentially. Finally, pH was adjusted to 7 with 1 mL of glacial acetic adic and the total volume adjusted to 1L. The medium was then aliquoted in glass bottles and autoclaved. 1 mM EGTA was added immediately prior to TIRF imaging. TAP-Ca is nominally Ca^2+^ free (free Ca^2+^<0.1 nM).

### TIRF microscopy

All imaging was performed on an inverted TIRF microscope (IX71, Olympus, Japan with an Olympus UApochromat 150x/1.45 Oil-immersion TIRFM objective) equipped with an EMCCD camera (iXon Ultra 897 BV, Andor, Belfast, Northern Ireland). The microscope is equipped with a custom-built TIRF condenser with manual TIRF-angle adjustment. TIRF illumination is provided by a fiber coupled 491 nm diode-pumped solid state laser. The microscope is fitted with a water hydraulic micro-manipulator (WR-6, Narishige, Japan), which was used to hold and position the wedge. All imaged were recorded at 81 ms frame time and 512×512 pixels at a resolution of 0.107 µm px^-1^. In the case of cells treated with Lithium, 100 µL of 33 mM LiCl was added to 0.5 mL of cells directly on the glass slide on the TIRF microscope without mixing.

### Analysis of TIRF Image Data

Recorded TIRF images were opened in Fiji (***Schindelin et al., 2012***), a de-speckle filter applied, and the background subtracted with a rolling-ball filter with a radius of 20 pixels. To correct for cells not sitting still, the stacks were maximum-projected in groups in order to identify all possible locations of the cilia, which were then traced individually using splines. Thus, selected splines were used to create kymograms from the filtered and background-subtracted movies using the multi-kymograph plugin (https://imagej.net/Multi_Kymograph), using a line-width of 5 pixels. Resulting kymograms were aligned using SIFT-C registration and finally maximum-projected to result in a kymogram covering the entire temporal duration of the recording. Finished kymograms were analyzed with a python script using OpenCV-v4.1.2 (***Itseez, 2015***) to identifies anterograde and retrograde trains by template matching with 8 px × 8 px templates of lines at angles of ±35° to ±55°, respectively. Resulting maps were curated for false positives and used to give false colors the original kymogram data.

### Cryo electron tomography

Whole *C. reinhardtii* cells (mat3-4, CC3994) were frozen by applying a light cell suspension directly onto the front side of glow-discharged holey carbon grids (Q350AR-35, Quantifoil Micro Tools, Germany). Cells were pretreated with 5 mM LiCl to increase the length of flagella. 10 nm gold beads were added in suspension as fiducial markers and the grids blotted from the back side before plunge freezing into liquid ethane.

Grids were imaged at 300kV on an FEI Titan Halo equipped with a field-emission gun, an imaging energy filter operated at 20eV and a K2 direct detection camera (both Gatan, USA). Cilia were located on full-grid low-magnification images and imaged bi-directionaly in 2 deg +ree steps from −20° to 60° and −22° to −60° in SerialEM (***Mastronarde, 2005***). Images were recorded at a nominal magnification of 30 000 in dose-fractioned super-resolution stacks of 10 images per tilt angle with a total of 3.2 e^-^ Å^-2^, resulting in a nominal pixel size of 2.35 Å px^-1^.

The tilt-series data was motion-corrected and binned once with MotionCor2-1.2.6 (***Zheng et al., 2017***). Resulting tilt-stacks were aligned, CTF-corrected in Etomo in IMOD-4.10.32 (***Mastronarde and Held, 2017***). To denoise the data, raw frame stacks were split into motion-corrected even/odd tilt frames. The previously generated alignment was applied to both even and odd stacks. These fine-aligned stacks were reconstructed into tomograms with ICON-GPU-v1.2.9 (***Chen et al., 2017***) on cluster nodes with two NVidia M2090 Tesla GPUs. Finally, the two tomograms were used as input to cryoCARE (***Buchholz et al., 2019a***,b) in order to generate a denoised tomogram.

## Acknowledgments

We would like to thank the Light Microscopy Facility (LMF) as well as the Electron Microscopy Facility (EMP) at MPI-CBG for their support. We would also like to thank Aliona Bogdanova from the Protein Expression and Purification (PEPC) Facility for cloning of the IFT46-mNeonGreen plasmid. Additionally we would like to thank Oscar Gonzales for assistance with high-performance computing resources. A.P.N is supported by an EMBO Long–term fellowship under ALTF number 891-2018 as well as by an HFSP Cross-disciplinary fellowship with reference number LT000515/2019. I.Z. was supported by the IMPRS Student Research Internship program. L.S. was supported by the Dresden International Graduate School for Biomedicine and Bioengineering (DIGS-BB), granted by the German Research Foundation (DFG) in the context of the Excellence Initiative. Finally, we would like to acknowledge the European Research Council (ERC) under the European Union’s Horizon 2020 research and innovation program (grant agreement No. 819826) to GP.

